# The geometry of habitat fragmentation: effects of distribution patterns on short-term species persistence

**DOI:** 10.1101/442467

**Authors:** Felix May, Benjamin Rosenbaum, Frank M. Schurr, Jonathan M. Chase

**Author notes:** Contact details: Felix May.

## Abstract

Land-use changes cause habitat loss and fragmentation and are thus important drivers of anthropogenic biodiversity change. However, there is an ongoing debate about how fragmentation *per se* affects biodiversity in a given amount of habitat. We illustrate why it is important to distinguish two different aspects of fragmentation to resolve this debate: (i) geometric fragmentation effects, which exclusively arise from the spatial distributions of species and habitat fragments, and (ii) demographic fragmentation effects due to reduced fragment size, increased isolation, or edge effects. While most empirical studies are primarily interested in quantifying demographic fragmentation effects, geometric effects are typically invoked only as post-hoc explanations of biodiversity responses to fragmentation *per se*. Here, we present an approach to quantify geometric fragmentation effects on species persistence probability. We illustrate this approach using spatial simulations where we systematically varied the initial abundances and distribution patterns (i.e. random, aggregated, and regular) of species as well as habitat amount and fragmentation *per se.* As expected, we found no geometric fragmentation effects when species were randomly distributed. However, when species were aggregated, we found positive effects of fragmentation *per se* on persistence probability for a large range of scenarios. For regular species distributions, we found weakly negative geometric effects. These findings are independent of the ecological mechanisms which generate non-random species distributions. Our study helps to reconcile seemingly contradictory results of previous fragmentation studies. Since intraspecific aggregation is a ubiquitous pattern in nature, our findings imply widespread positive geometric fragmentation effects. This expectation is supported by many studies that find positive effects of fragmentation *per se* on species occurrences and diversity after controlling for habitat amount. We outline how to disentangle geometric and demographic effects of fragmentation, which is critical for predicting the response of biodiversity to landscape change.

## Introduction

Anthropogenic land-use changes cause the loss, the fragmentation, and the degradation of natural and semi-natural habitats (Harrison and Bruna 1999, Fischer and Lindenmayer 2007) and are considered as one of the most important drivers of past, current, and future biodiversity change (Millennium Ecosystem Assessment 2005, Pereira et al. 2010, Pimm et al. 2014, Newbold et al. 2015). Each of these three processes – habitat loss, fragmentation, and degradation – interact to alter biodiversity in the face of anthropogenic pressures, but because they often act in concert, it is difficult to disentangle their influences (Didham et al. 2012).

The concept of fragmentation, in particular, has generated a lot of debate and confusion. This is because the term ‘fragmentation’ is referred to both as a dynamic process (i.e., change in a given landscape through time from continuous to fragmented natural habitat) and as a static pattern (i.e., some landscapes have higher degrees of fragmentation than others) (Fahrig 2003, 2017). From the dynamic perspective, fragmentation involves a reduction in habitat amount (i.e. habitat loss), as well as a changes in spatial habitat configuration (Didham et al. 2012). From the static pattern-based perspective, fragmentation – also called fragmentation *per se* to avoid ambiguity (Fahrig 2003) – refers to the spatial configuration of a constant amount of habitat at a given point in time. In this study, we are explicitly interested in disentangling the independent consequences of habitat amount and fragmentation *per se*, and thus focus on landscapes with different static configurations of a given habitat amount. Indeed, understanding and predicting the distinct consequences of habitat amount and fragmentation *per se* is important to evaluate alternative spatial scenarios of landscape change for conservation and land-use management (Tscharntke et al. 2012).

The issue of the consequences of fragmentation *per se* independent of habitat amount is closely related to the question of whether it is better for biodiversity conservation to preserve a single large (SL) or several small (SS) habitat fragments (Fahrig 2013). The latter is well known as the SLOSS problem in conservation biology (e.g. Diamond 1975, Simberloff and Abele 1976, Ovaskainen 2002, Tjørve 2010), which remains unresolved even after four decades of research. Although numerous studies have shown the often expected negative effects of fragmentation *per se* on biodiversity (e.g. Rybicki and Hanski 2013, Haddad et al. 2015), a great many find neutral (Fahrig 2003, 2013, Yaacobi et al. 2007) or even positive effects (e.g. Tscharntke et al. 2002, Fahrig 2017, Seibold et al. 2017).

To understand and resolve the contrasting results observed in empirical studies, we suggest distinguishing two different aspects of fragmentation: (i) geometric fragmentation effects, which arise solely from the spatial arrangement of habitat fragments relative to species distributions in continuous landscapes and specify if individuals are located in habitat fragments or in the surrounding (hostile) matrix, and (ii) demographic fragmentation effects, which alter population growth or persistence in potentially isolated habitat fragments due to mechanisms such as increased demographic stochasticity (MacArthur and Wilson 1967, Lande 1993), Allee effects (Courchamp et al. 2008, Swift and Hannon 2010), reduced immigration (Hanski 1999, Hanski et al. 2013), or edge effects (Saunders et al. 1991, Harrison and Bruna 1999, Collinge 2009, Haddad et al. 2015). While geometric effects by definition only depend on the spatial distributions of species and habitat fragments, demographic effects depend on species traits and their potentially complex interactions with the modified environment.

In this study, we focus on geometric fragmentation effects, although it is important to acknowledge that geometric and demographic effects can work at the same time and thus will need to be simultaneously considered in empirical studies. Geometric fragmentation effects have been well-known for several decades (Diamond 1975, Quinn and Harrison 1988) and they are often qualitatively discussed as post-hoc explanations of observed fragmentation *per se*-biodiversity relationships (e.g. Tscharntke et al. 2002, Seibold et al. 2017). However, while these effects are known conceptually, we still require tools to quantify and predict this important aspect of fragmentation (Raheem et al. 2009, Tscharntke et al. 2012).

When only geometric effects are considered, it is assumed that habitat fragments work like a cookie-cutter, which means all individuals survive in habitat fragments, but die in the matrix. This simplified perspective intentionally ignores more complex biological responses in space and time. For the purpose of this study, we also define persistence and extinction at the landscape-scale based exclusively on the geometry of species’ and fragments’ distributions. Landscape-scale extinction means that all individuals of a given species are located in the matrix, while landscape-scale species persistence means that one or more individuals of the focal species are located in habitat fragments (Fig. 1). Accordingly, this geometric definition of species persistence purposely excludes long-term responses due to demographic fragmentation effects (Kuussaari et al. 2009, May et al. 2013) and/or the ability of species to persist in the matrix (Pereira and Daily 2006, Fischer and Lindenmayer 2006, Didham et al. 2012).

**Figure 1:**
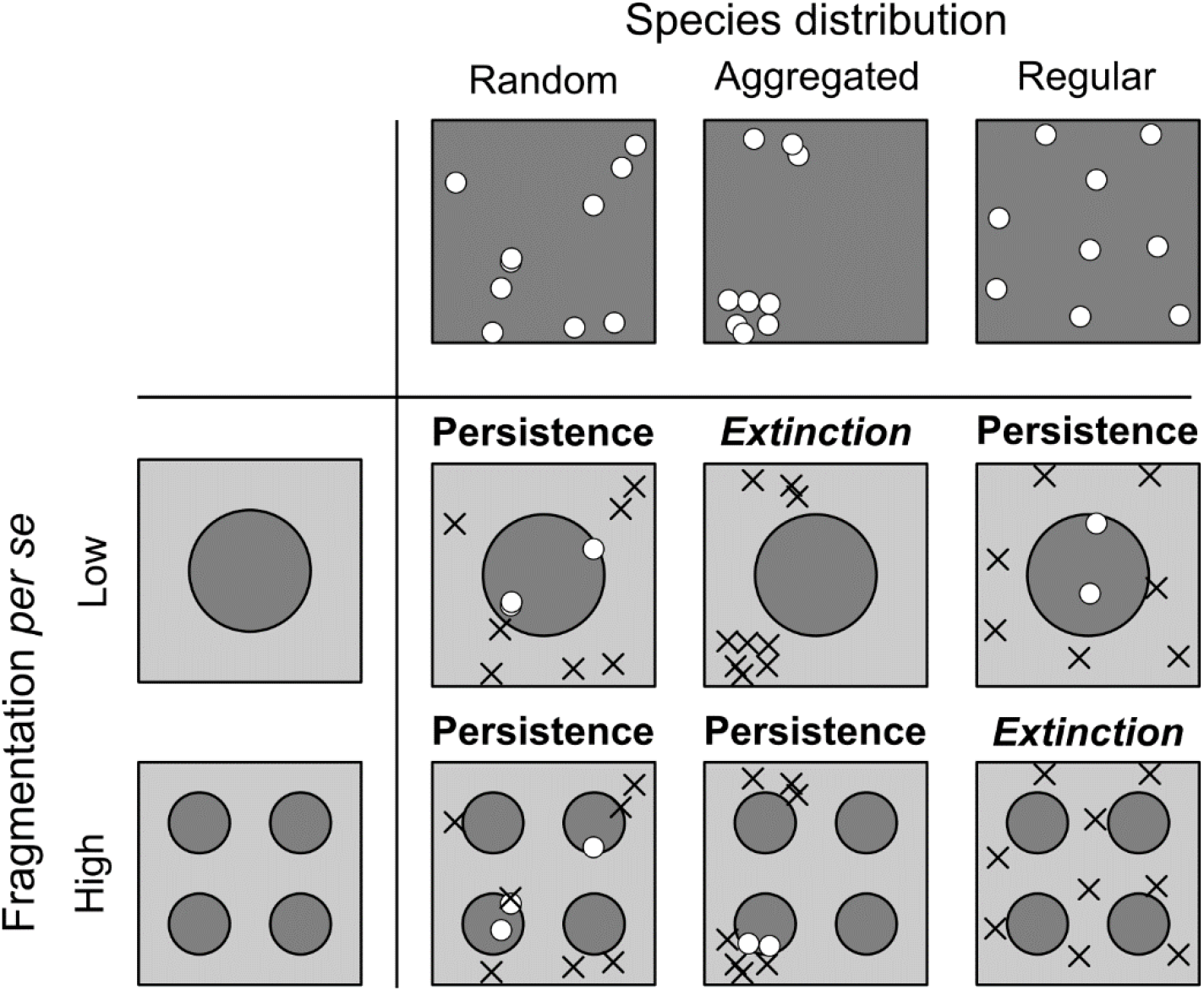
Illustration of geometric fragmentation effects on landscape-scale species persistence. Dark grey areas indicate suitable habitat, while light grey areas indicate unsuitable matrix, where individuals cannot survive. Accordingly, white points indicate living individuals, while black crosses indicate individuals that died in the matrix. In case of random species distributions persistence probability is independent of fragmentation. With aggregated species distributions, however, species persistence probability (i.e. at least one individual in habitat) increases with fragmentation *per se*. With regular distributions, persistence probability might decrease with fragmentation *per se*, because fragments fit into the gaps between individuals. The interactive effects between species distributions, habitat amount, and fragmentation *per se* on abundances and persistence probabilities are quantified in detail in this study.

Geometric fragmentation effects depend on the distributions of individuals and species in continuous habitat prior to landscape change. When the distribution of individuals follows complete spatial randomness (CSR), the number of individuals in the remaining fragments only depends on the total habitat amount. Under a CSR distribution, we thus expect landscape-scale species persistence, as defined above, to depend on habitat amount, but not on fragmentation *per se* (Fig. 1, left column). In contrast, when species show intraspecific aggregation, where individuals of the same species occur closer together than expected in a CSR distribution, several smaller fragments are more likely to “sample” at least a few individuals than a single large fragment (Fig. 1, middle column). Therefore, we might expect fragmentation *per se* to be beneficial for the persistence of species with aggregated, i.e. clustered, distributions. When species show a regular distribution of their individuals, all individuals are separated by similar distances and there are gaps with a typical size among the individuals. In this case, small fragments might be located in the gaps between species, while a large fragment, which is larger than the gap size, will include at least some individuals. Accordingly, we expect persistence to decrease with fragmentation *per se* (Fig. 1, right column).

It is important to note that geometric fragmentation effects occur regardless of the mechanism that generates non-random species distributions (i.e. aggregation or regularity) in continuous habitat. In landscapes with pre-existing environmental heterogeneity, species will often show aggregated distributions, because of their species-specific habitat requirements and changes of competitive ability with environmental conditions (Whittaker 1962, 1975, Chase and Leibold 2003). However, non-random species distributions can emerge even without environmental heterogeneity, when intraspecific aggregation is generated by processes such as dispersal limitation (Hubbell 2001, Chave et al. 2002) and/or positive density dependence (Courchamp et al. 1999, Molofsky et al. 2001), while regular distributions can be generated by negative density dependence, due to resource competition and/or species-specific pathogens or consumers (Sterner et al. 1986, Bagchi et al. 2011). In nature, aggregated species distributions appear to be the rule rather than the exception and may be among the most fundamental patterns in ecology (Condit et al. 2000, McGill 2010, 2011).

Geometric fragmentation effects have rarely been explicitly studied because theoreticians and empiricists have mostly focussed on demographic fragmentation effects, while geometric effects are often qualitatively invoked, usually with references to habitat heterogeneity and the resulting beta-diversity, to explain observed positive relationships between fragmentation per se and biodiversity from a post-hoc perspective (reviews in Quinn and Harrison 1988, Fahrig 2017). Moreover, we still lack approaches for the quantification of geometric effects. Recently, Chisholm et al. (2018) presented an approach for quantification of geometric effects, called short-term species loss in their study. However, their approach focusses on species distributions predicted by neutral models, which by definition exclude regular distributions and assume that all species share the same dispersal parameters and thus show similar spatial distributions (Hubbell 2001). Here, we address these gaps by presenting a more generic approach to quantify geometric fragmentation effects on species abundances and their persistence. This approach allows us to efficiently evaluate a large range of scenarios with respect to species abundances and distributions prior to landscape change, as well as a large range of landscape configuration scenarios, including variations in habitat amount and fragmentation *per se*. The simulated species distributions include random, aggregated, and regular distributions. We will show that species mean abundances are only determined by habitat amount, but not by fragmentation *per se*. In contrast, the effect of fragmentation *per se* on persistence depends on the spatial pattern of the species distribution. For random distributions, there is no effect of fragmentation *per se* on persistence probability, while the effects is consistently positive for aggregated distributions and weakly negative for regular species distributions. We argue that it is essential to understand the consequences of both geometric and demographic fragmentation effects in order to reconcile the mixed results of previous research and to advance the debate about the consequences of fragmentation for biodiversity.

## Materials and Methods

In order to quantify geometric fragmentation effects, we develop a simulation approach that predicts species abundances and persistence probabilities in fragmented landscapes according to the geometric definition provided above. We designed the simulations in a way that enables one to independently vary habitat amount and fragmentation *per se*. Furthermore, we can manipulate species abundances and their spatial distributions in continuous landscapes prior to landscape change. Simulated species distributions include random, aggregated and regular patterns (Fig. 1).

### Species distributions

We simulate species distributions using point process models, where every point represents one individual (Wiegand and Moloney 2014, Baddeley et al. 2015). For random distributions, we use the Poisson process, which assumes complete spatial randomness (CSR) without any interactions between individuals in the simulated arena. We simulated random distributions with a density of 10, 100, and 1,000 points in a square arena of 1 × 1 units.

We model aggregated species distributions using the Thomas process, which is a special case of the Poisson cluster process (Thomas 1949, Morlon et al. 2008, Wiegand and Moloney 2014). The aggregated distribution of individuals are defined by the following steps (Morlon et al. 2008, Afshang et al. 2017):

1. The centres of clusters are distributed according to a Poisson process (i.e. complete spatial randomness – CSR), in a square landscape with area *A*. The density of the cluster centres is given by ρ.
2. The numbers of individuals in each cluster is randomly assigned from a Poisson distribution with mean and variance μ.
3. The positions of individuals around the cluster centre is modelled using a bivariate radially symmetric Gaussian distribution with mean 0 and variance σ^2^.

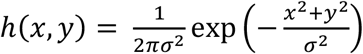

According to these simple rules the expected number of clusters is given by n_c_ = ρ*A, the average number of individuals per cluster is μ and the spatial extent of the cluster is associated with σ. Due to the random distribution of cluster centres and numbers of points per cluster, clusters can overlap and/or there can be empty clusters with zero individuals. Therefore, the real number of clusters can deviate from n_C_. The expected total abundance (i.e. the number of points) of the species is n_P_ = ρ*A*μ. We used the function `sim_thomas_community` from the R package `mobsim` to simulate the species distributions following the Thomas process (May et al. 2018). We simulated a full factorial design of the parameter combinations n_P_ in {10; 100; 1,000}, n_C_ in {1; 2; 5; 10}, and σ in {0.01, 0.02, 0.05, 0.1}.

For simulations of regular species distributions, we applied the Strauss process, which combines inhibition of individuals at small spatial scales with randomness at larger scales (Sterner et al. 1986, Wiegand and Moloney 2014). In the Strauss process there is inhibition of neighbouring individuals at distances smaller than the interactions radius r. The strength of the inhibition is governed by parameter γ, which is in the interval [0, 1]. When γ = 0, there is perfect inhibition, which means the minimum distance between points is expected to be r. When γ = 1 there is no inhibition anymore and the Strauss process converges to the Poisson process (CSR). In the simulations we defined the inhibition strength (γ) as a free parameter, but we derived the inhibition distance from the simulated number of individuals (N) as r = 1/√N, which is the distance between individuals if they are arranged in a perfect lattice that covers that total landscape area. We simulated realizations of the Strauss process using the Metropolis-Hastings algorithm as implemented in the function `rmh` in the R package `spatstat` (Baddeley et al. 2015). Again, we conducted simulations for a full factorial design of the parameter values n_P_ in {10; 100; 1,000} and γ in {0, 0.01, 0.1, 1}.

### Fragmented landscapes

We simulated fractal raster maps with defined habitat amount and fragmentation *per se* (With 1997, With et al. 1997, Körner and Jeltsch 2008, Campos et al. 2013). For this purpose we used the midpoint displacement algorithm (Saupe 1988) as implemented in the R package `FieldSim`. This algorithm generates 3-dimensional fractal surfaces, where the ruggedness of the surface is controlled by the Hurst factor (H). This parameter is defined in the interval from 0 (rugged surface) to 1 (smooth surface) and is related to the fractal dimension D of the surface by D = 3.0 – H (Saupe 1988). Slicing the surface at a given “elevation” allows to define the habitat amount of the landscape (Fig. 2). All raster cells above a certain threshold are defined as habitat and all cells below the threshold as matrix. The Hurst factor (H) then defines the fragmentation *per se* of the landscape, where varying H from 0 to 1 represents landscapes from high to low fragmentation *per se* (With 1997, Campos et al. 2013) (Fig. 2). We generated raster maps of 129 × 129 grid cells and varied habitat amount – measured as proportion of habitat in the landscape – from 0.01 to 0.5 and fragmentation *per se* from H = 0.9 (low fragmentation *per se*) to 0.1 (high fragmentation *per se*).

**Figure 2:**
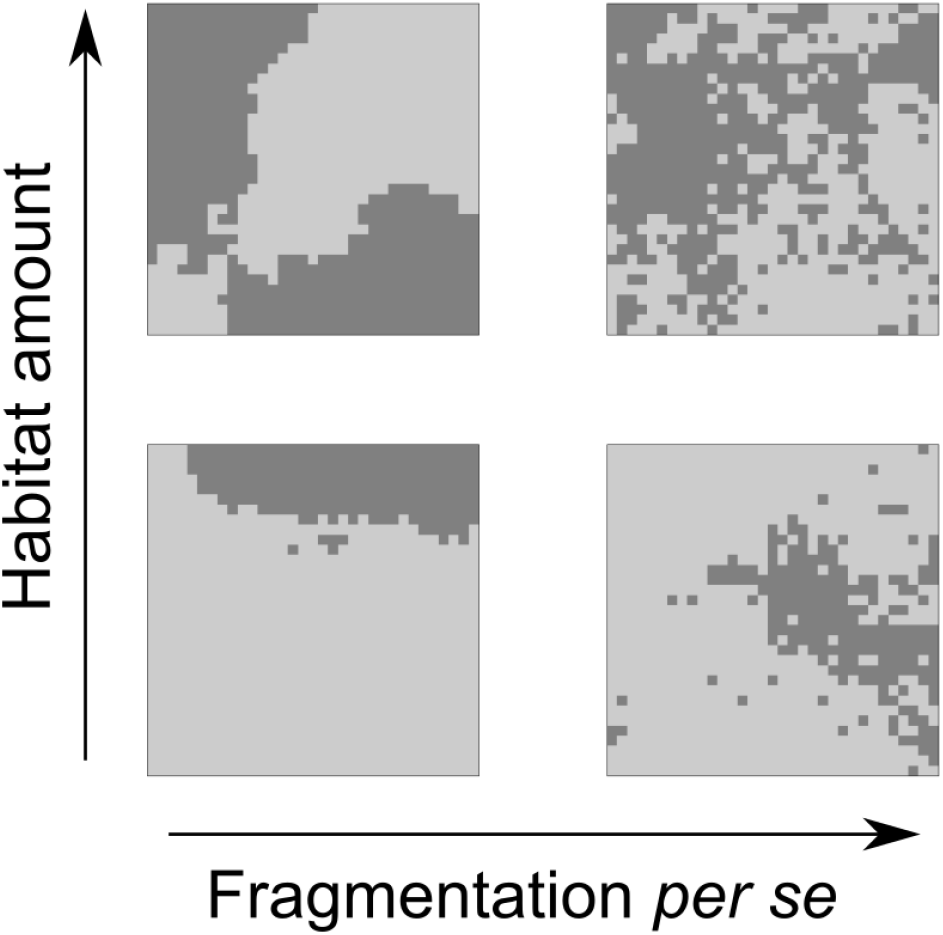
Fractal landscapes simulate with the midpoint displacement algorithm. Dark grey areas indicate suitable habitat, while light grey areas indicate unsuitable matrix. The landscapes shown here consist of 33 × 33 grid cells for better visualization, while 129 × 129 cells were used for the simulations of species abundances and persistence. The top row and bottom rows show landscapes with 50% and 20% of habitat. The left and right columns show landscapes with Hurst factor (H) of 0.9 (low fragmentation *per se*) and 0.1 (high fragmentation *per se*).

### Species abundances and persistence in fragmented landscapes

Species abundances and persistence in fragmented landscapes were evaluated by simply overlaying point patterns representing the species distribution in continuous landscapes and the fractal maps representing the habitat distribution in fragmented landscapes. All individuals in habitat cells were labelled as survivors and all individuals in the matrix were removed (Fig. 1). For each scenario of initial species abundance, distribution and habitat fragmentation, we conducted 1,000 replicate simulations and recorded the species abundance and persistence (i.e. abundance > 0) in the fragmented landscape.

For aggregated species distributions modelled by the Thomas process and the special case of landscapes with equally-sized circular habitat fragments there is an analytical solution for the persistence probability (Afshang et al. 2017) (see Appendix 1). We compare results from this analytical solution with our stochastic simulation approach in Appendix 2 (Figs. A2, A3).

## Results

For all simulation scenarios of initial species abundances, distributions, habitat amount, and fragmentation *per se* we evaluated the mean and variation of abundance in fragmented landscapes, as well as the geometrically defined persistence probability based on 1,000 replicate simulations. In a first step, we compared responses to habitat amount and fragmentation *per se* among the three distribution types random, aggregated, and regular. In subsequent steps, we investigated how persistence probabilities in fragmented landscapes vary with respect to specific parameters of species abundances and their distributions.

By statistical definition the expected number of individuals in habitat fragments equals the initial abundance times the proportion of habitat (i.e. habitat amount) independent of the spatial configuration of habitat. Accordingly, in our simulations the mean abundance across replicates was always independent of fragmentation *per se* and equalled the initial abundance (n_P_) times the habitat amount (Fig. 3, top row). However, fragmentation *per se* clearly influenced the variability of abundance among replicate simulations and this relationship qualitatively and quantitatively changed with species distribution patterns (Fig. 3, middle row). With random distributions (CSR) the coefficient of variation (cv) of abundances was independent of fragmentation *per se*. With aggregated distributions variation was always higher than with CSR and furthermore variation decreases with increasing fragmentation *per se*. For regular distributions, we found the opposite results that means variation was consistently lower than with CSR and slightly increases with fragmentation *per se*.

**Figure 3:**
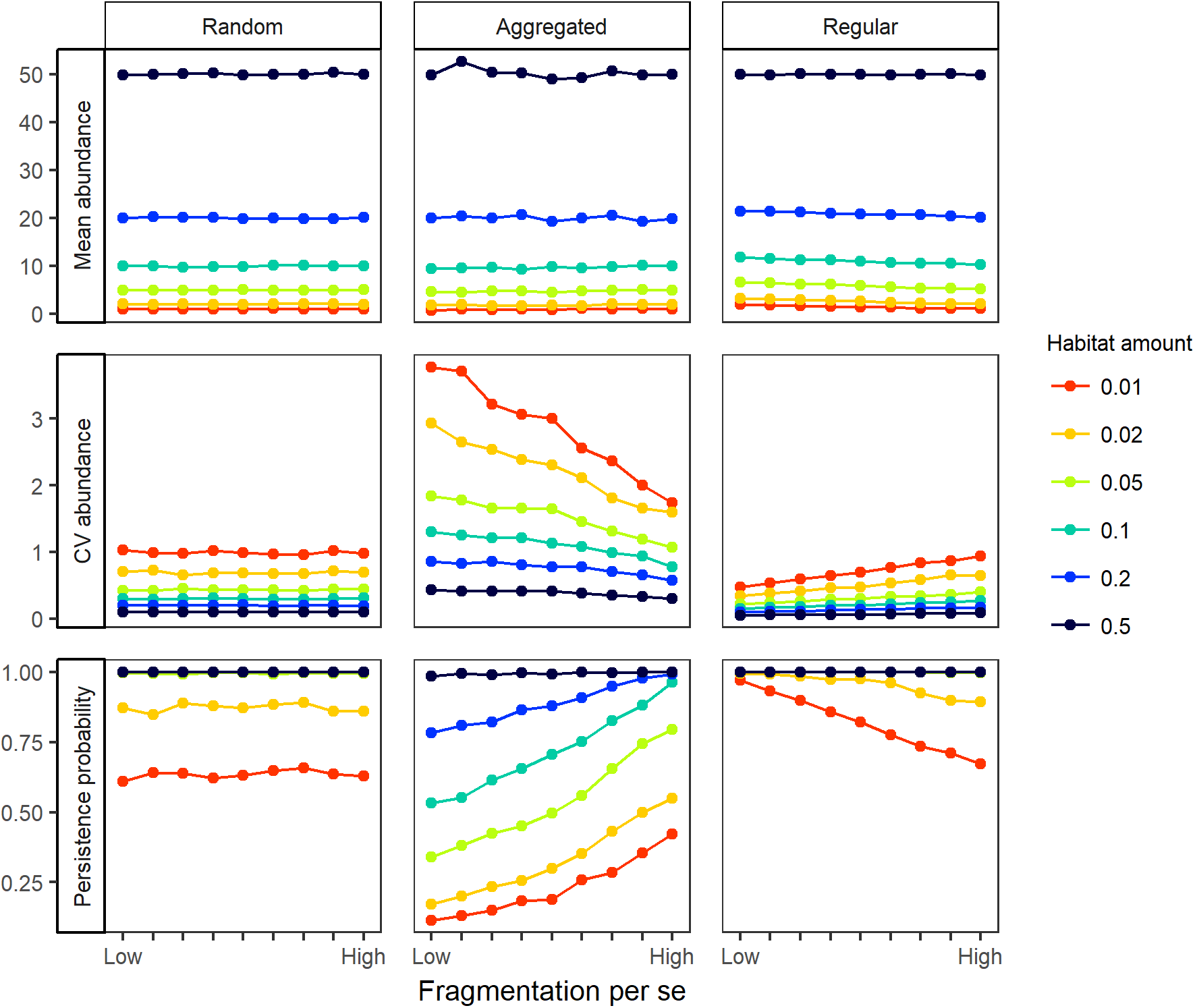
Consequences of habitat amount and fragmentation *per se* for species abundance and persistence with random, aggregated and regular species distributions. All simulations were conducted with 100 individuals. In the aggregated scenario (middle column) we used a cluster size of σ = 0.02 and a number of clusters of n_C_ = 5. In the regular scenario (right column), we used an inhibition strength parameter of γ = 0.01. The rows show the mean abundance (top), coefficient of variation of abundance (= standard deviation/mean) (middle) and the persistence probability (bottom) estimated from 1,000 replicate simulations. Low fragmentation corresponds to a Hurst factor (H) of 0.9 and high fragmentation per se to H = 0.1. The line colour indicates the habitat amount measured as proportion of habitat in the landscape.

These changes in variation of abundance also resulted in changes in persistence probability, i.e. the probability that abundance is larger than zero in the fragmented landscape (Fig. 3, bottom row). Of course, persistence probability consistently increased with increasing species abundances (results not shown). With respect to species distributions, persistence probability was highest with regular distributions, intermediate with random and lowest with aggregated distributions. Persistence probability was independent of fragmentation *per se* with random distributions. With intraspecific aggregation persistence probability increased with fragmentation *per se*, but this increase was also influenced by habitat amount. The positive relationship of persistence probability to fragmentation *per se* was strong for low or intermediate habitat amounts, but disappeared for high habitat amounts. In contrast, with regular distributions persistence probability slightly decreased with fragmentation *per se*, but only for low habitat amount. The effect of fragmentation *per se* was weaker with regular compared to aggregated distributions.

In a second step, we assessed the consequences of species distribution parameters within aggregated and regular species distributions on persistence probabilities in more detail. The overall aggregation of a species whose distribution follows the Thomas process depends on the number of clusters (n_C_), the size of clusters (σ), and the number of individuals per cluster (μ). The total abundance (n_P_) is the product of number of clusters (n_C_) and the number of individuals per cluster (μ). Therefore, just two out of these three parameters (n_P_, n_C_, μ) can be varied independently, while the cluster size parameter (σ) is independent of all the others. We analysed how persistence probability in fragmented landscapes varies with changing cluster size and number of clusters for species with fixed total abundance (100 individuals). We found that persistence probability increased with the number of clusters and with the size of clusters, in agreement with the general result that the persistence probability is higher for less aggregated species distributions (Fig. 4). In almost all scenarios there was an increase in persistence probability with fragmentation *per se*. The positive effect of many small fragments only vanished for high numbers of clusters, large cluster size, and high habitat amount (Fig. 4, lower right corner).

**Figure 4:**
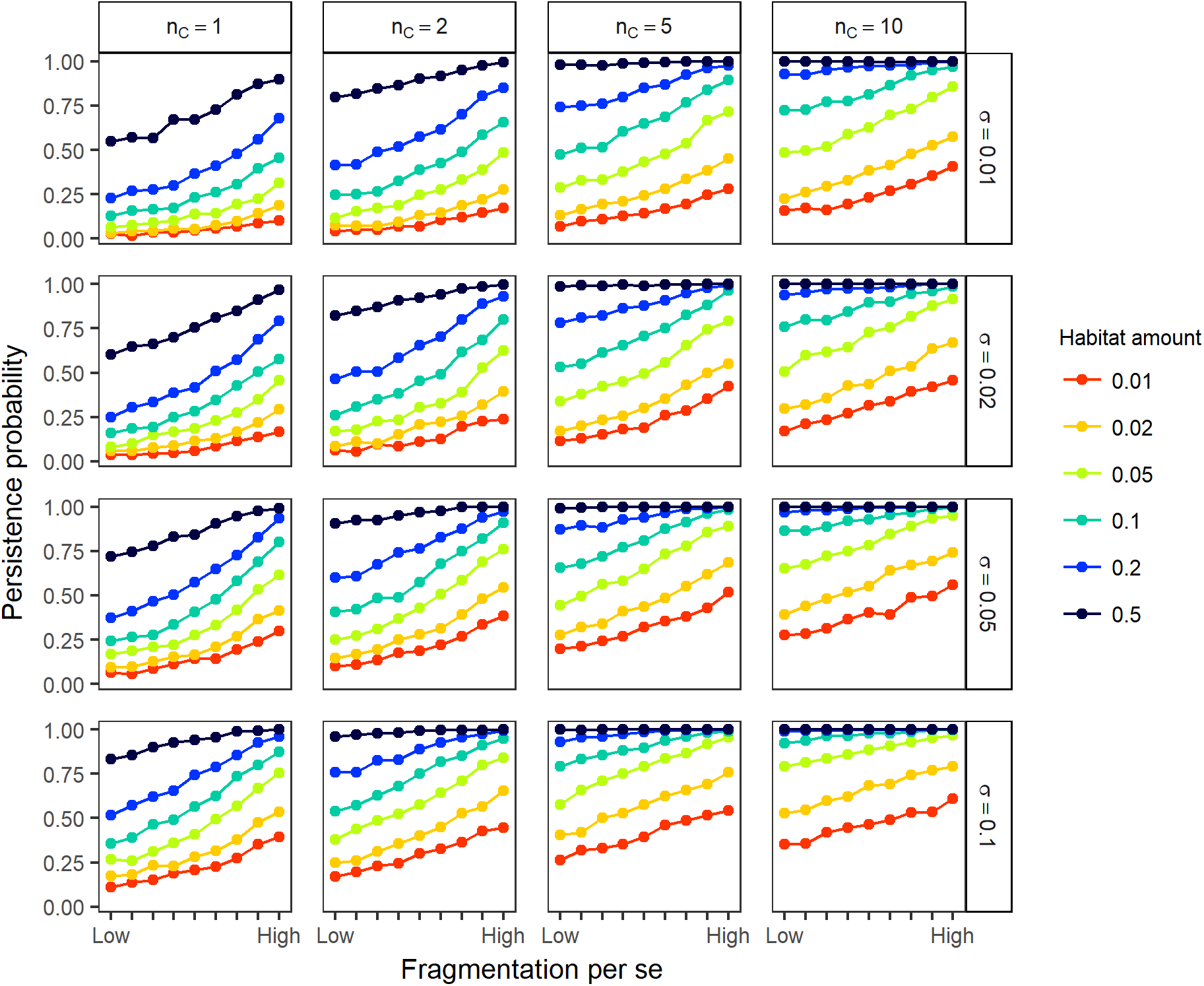
Consequences of habitat amount and fragmentation *per se* for species persistence probability with aggregated species distributions. All simulations were conducted with 100 individuals. The rows and columns show different aggregation scenarios with different cluster sizes (σ) in rows and different numbers of clusters (n_C_) in columns. Persistence probabilities were estimated from 1,000 replicate simulations. Low fragmentation *per se* corresponds to a Hurst factor (H) of 0.9 and high fragmentation *per se* to H = 0.1. The line colour indicates the habitat amount measured as proportion of habitat in the landscape.

For regular distributions, we assessed how the effect of fragmentation *per se* on persistence probability changed with the strength of neighbour inhibition (γ). The negative relationship between fragmentation *per se* and persistence probability was stronger for stronger inhibition (i.e. lower γ-values that increase regularity, Fig. 5). The negative effect of fragmentation *per se* also vanished with increasing habitat amount. For landscapes with more than 5% of habitat amount there were only effects of fragmentation *per se* on persistence for the very low initial abundances of 10 individuals (Fig. 5, bottom).

**Figure 5:**
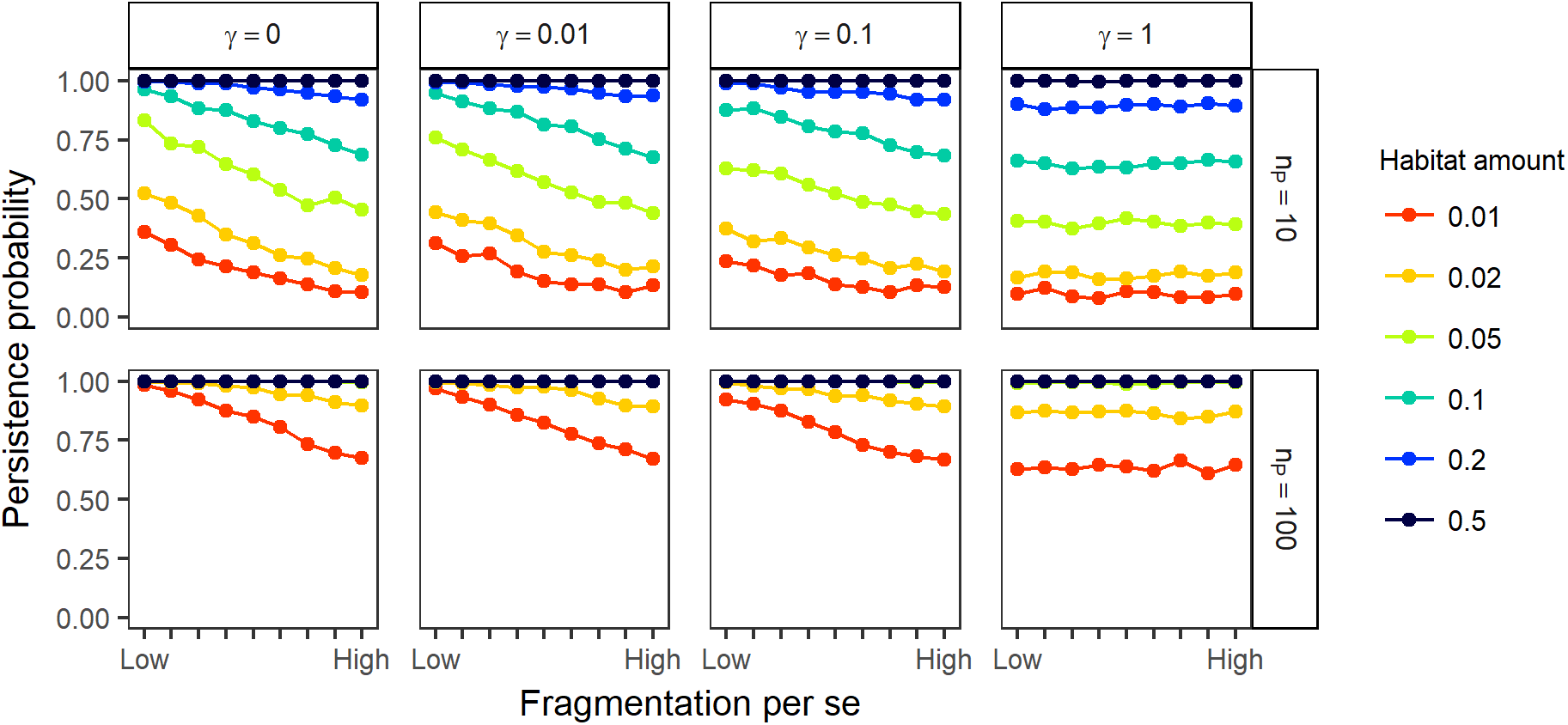
Consequences of habitat amount and fragmentation *per se* for species persistence probability with regular species distributions. The rows and columns show different distribution scenarios with different abundances prior to fragmentation (n_P_) in rows and different strength of neighbour inhibition (γ) in columns. Persistence probabilities were estimated from 1,000 replicate simulations. Please note that γ = 1 corresponds to complete spatial randomness (CSR). Low fragmentation *per se* corresponds to a Hurst factor (H) of 0.9 and high fragmentation *per se* to H = 0.1. The line colour indicates the habitat amount measured as proportion of habitat in the landscape.

All of these simulation results made use of habitat maps generated with the midpoint displacement algorithm for fractal landscapes. However, for landscapes where we assume equally-sized circular fragments, which are randomly distributed, there is an analytical solution for the persistence probability for aggregated distributions modelled with the Thomas process (see Appendix 1). This analytical solution was in agreement with our simulation results, indicating that the results reported here do not depend on idiosyncrasies of the fractal habitat maps used (Appendices 1 and 2, Figs. A2, A3, A4).

## Discussion

The consequences of habitat loss and fragmentation *per se* for species persistence and biodiversity are driven by two distinct consequences of landscape change. First, there are geometric fragmentation effects, which arise from the distribution of individuals inside or outside of habitat fragments. Second, there can be demographic responses to fragmentation, such as increased demographic stochasticity, Allee effects, reduced immigration, or edge effects in habitat fragments (e.g. Hanski 1999, Courchamp et al. 2008, Collinge 2009). We argue that much confusion about fragmentation has been arising because geometric effects have either been ignored or not clearly distinguished from demographic effects (e.g. Hanski et al. 2013, Haddad et al. 2015).

In this study, we present an approach to quantify geometric fragmentation effects. Specifically, our approach predicts species short-term persistence probability as function of species abundances and distributions prior to landscape change, and as function of habitat amount and fragmentation *per se*. We found consistent positive geometric fragmentation effects of fragmentation *per se* on persistence when species had aggregated distributions and weakly negative geometric effects when species had regular distributions. While especially the first finding has been reported qualitatively before (e.g. Quinn and Harrison 1988, Tscharntke et al. 2002, Seibold et al. 2017), our approach allows quantifying the strength of geometric fragmentation effects for a large range of species and habitat distribution scenarios.

The distinction between geometric and demographic fragmentation effects sheds new light on previous studies of fragmentation effects on biodiversity. Recently, Fahrig (2017) reviewed 118 studies that found significant effects of fragmentation *per se* on species richness, species abundances, or occurrences, while controlling for habitat amount. Out of the significant effects of fragmentation *per se*, 76% were positive and 24% were negative. Similar results have been also found in previous reviews of the SLOSS problem (Simberloff and Abele 1982, Quinn and Harrison 1988). From the perspective adopted in our study, these findings might indicate that often positive geometric fragmentation effects outweigh the presumably mostly negative demographic effects of fragmentation *per se*. The review of Fahrig (2017) only considers studies which reported significant fragmentation effects. However, there are also many studies that did not find clear effects of fragmentation *per se* (Yaacobi et al. 2007, e.g. Fahrig 2013). Our findings offer an interesting novel interpretation of these studies. We speculate that in these studies, positive geometric effects might balance negative demographic effects so that the net effect cannot be detected statistically. Of course all these interpretations remain speculative for two main reasons. First, because we often lack information on species distributions and abundances in the continuous landscape prior to landscape change, and second, because there might be also positive demographic effects of fragmentation for specific taxa or trophic levels (reviewed in Fahrig 2017). For instance, fragmentation can prevent the spread of focal species’ antagonists such as competitors, herbivores, predators, or pathogens (e.g. Crooks and Soule 1999, Schippers et al. 2015, Brudvig et al. 2015), or can foster species that benefit from habitat-matrix edges, whose total length increases with fragmentation *per se* (e.g. Klingbeil and Willig 2009, Barrera et al. 2015). However, the ambiguity of previous fragmentation studies underlines the urgent need to quantitatively disentangle geometric and demographic effects of fragmentation.

In real landscapes, the overall consequences of fragmentation *per se* will most likely include both geometric and demographic effects. Our findings highlight the need that studies interested in demographic fragmentation effects should incorporate geometric effects as a null-hypothesis. According to our results and due to the generality of non-random – especially aggregated – species distributions in nature, the null-hypothesis of no significant geometric fragmentation effects is likely to be inappropriate. We suggest that depending on the degree of intraspecific aggregation prior to landscape change, a positive relationship between fragmentation and species persistence or biodiversity, respectively, is a more realistic null-hypothesis.

While our study exclusively investigated geometric effects, our findings suggest a protocol for separating geometric and demographic fragmentation effects based on species distribution data in modified landscapes. First, a reference scenario needs to be derived that is based only on geometric effects. Depending on the available data this can be done in different ways. When the data includes both, observations from a modified landscape and from an unmodified reference landscape with continuous natural habitat (e.g. Schmiegelow et al. 1997, Laurance et al. 2002), geometric fragmentation effects can be simulated in the continuous landscape using habitat distributions that represent the habitat amount and fragmentation *per se* in the observed modified landscape. This procedure is equivalent to the simulations we used here, except that observed species distributions are used instead of simulated ones. Unfortunately, there will often be no data from a continuous control landscape (e.g. Giladi et al. 2011). In this case it might still be possible to model species distributions by fitting a spatial model (e.g. the Thomas process or the Strauss process) to species distribution data from the largest habitat fragments available in the data set. This field-parameterized spatial model can then be used to simulate expected species distributions in continuous habitat prior to landscape change (e.g. Plotkin et al. 2000, Morlon et al. 2008) and to estimate geometric fragmentation effects as suggested in the first approach. Both of these approaches estimate pure geometric fragmentation effects, irrespective of demographic changes. The second step is then to compare observed species distributions or biodiversity data from fragmented landscapes to the null expectation representing only geometric effects. The difference between the null expectation and the field observations provides an estimate for the demographic consequences of fragmentation with appropriate control for geometric effects.

In this study, we used specific spatial models of species aggregation (the Thomas process) and regularity (the Strauss process). These models allow varying different components of species distributions, specifically the number and sizes of clusters as well as the numbers of individuals per cluster in the Thomas process, and the strength of neighbour inhibition in the Strauss process. We investigated all of these components systematically and in combination with two different approaches for simulations of habitat distributions, specifically fractal landscapes and landscapes with and equally-sized circular fragments. Therefore, we are confident that our results do not depend on a specific model. We consider it as an advantage of our study that aggregation and regularity are modelled in a generic way without reference to a specific ecological mechanism. This means that our findings encompass ecosystems where non-random distributions can be caused by distinct processes, including environmental heterogeneity plus habitat filtering, local dispersal, competition, or facilitation among conspecific individuals. Several previous studies used spatially-implicit models, such as species-area relationships (Harte and Kinzig 1997, Kinzig and Harte 2000) or the negative binomials distribution (He and Legendre 2002, Green and Ostling 2003), to describe spatial distributions of species. Due to these spatially-implicit approaches these studies could only asses the consequences of habitat amount, but not of fragmentation *per se* for species diversity, which requires a spatially-explicit approach as used in this study.

Tjørve (2010) applied species-area curves in order to resolve different results within the SLOSS debate. Based on this approach he suggested that increasing species aggregation within fragments favours low fragmentation *per se* (i.e., a single large fragment), which seems to be in contrast with our finding that higher fragmentation *per se* maximizes persistence probability with aggregation. At the same time Tjørve (2010) predicted that decreasing overlap between fragments (i.e., higher beta-diversity) favours higher fragmentation *per se*, (several small fragments). However, while the independent variation of aggregation within fragments and beta-diversity among fragments is possible with a phenomenological approach such as species-area relationships, these two parameters will be coupled from a more mechanistic metacommunity perspective (Hubbell 2001, Condit et al. 2002). Increasing aggregation will usually increase beta-diversity and thus reduce the overlap among fragments, which in turn favours the several small strategy and high fragmentation *per se* according to our as well as Tjørve’s (2010) findings.

Dynamic and spatially-explicit simulation models offer an interesting approach to investigate the interplay between geometric and demographic fragmentation effects. Unfortunately, even models that potentially include both, geometric and demographic effects (e.g. Hanski et al. 2013, Rybicki and Hanski 2013) did rarely attempt to disentangle both aspects, but only focussed on the total effects of fragmentation on biodiversity. In this case it depends on model idiosyncrasies if the studies highlight overall negative (Rybicki and Hanski 2013) or overall positive (Campos et al. 2013) consequences of fragmentation *per se* with limited understanding of the relative contributions of geometric vs. demographic effects. Claudino et al. (2015) provided an attempt to disentangle geometric (called static) and demographic (called dynamic) fragmentation effects for communities simulated by a spatially-explicit neutral model. However, they do not address the question how to transfer their approach to non-neutral communities or to empirical data.

Important simplifying assumptions of the approach we use are the instantaneous and complete removal of all individuals outside of natural habitat fragments. In real landscapes, habitat transformation is not an instantaneous process, but happens over prolonged periods of time (de Barros Ferraz et al. 2005, Ewers et al. 2013, Claudino et al. 2015). Furthermore, species responses to landscape change can show time-lags (Kuussaari et al. 2009), and species might survive in the matrix and potentially disperse from the matrix to habitat fragments (Prevedello and Vieira 2010, Koh and Ghazoul 2010, Pereira et al. 2014). Investigating geometric and demographic fragmentation effects in systems with time-delayed responses and survival in the matrix is an important next step. In landscapes with rapid landscape change and high matrix mortality, temporal data could be used to disentangle geometric and demographic effects. While short-term responses of species and communities at the landscape-scale will primarily reflect geometric effects, long-term effects will be caused demographic changes (Kuussaari et al. 2009, Helm et al. 2009, Jones et al. 2016)

There has been a long and intense debate on the role of fragmentation for biodiversity. Our study highlights the need to distinguish and explicitly consider geometric and demographic fragmentation effects as a key issue for resolution of the debate. In retrospect, it is perhaps surprising that positive correlations of biodiversity with fragmentation *per se* are often perceived as unexpected (Fahrig 2017), despite well-established qualitative knowledge about positive geometric fragmentation effects. We hope that the approach and findings presented here will foster research on the relative importance of geometric and demographic fragmentation effects across taxa, ecosystems, and spatio-temporal scales, and provide a way forward for synthesizing seemingly contradictory results on responses of biodiversity to habitat loss and fragmentation *per se*.

## Data Accessibility Statement

This article is based on simulations and analytical calculations and does therefore not include any data.

## Authorship contributions

FM, FMS, and JMC designed the study. FM implemented and analysed the stochastic simulations and BR implemented the analytical calculations. FM wrote the first manuscript draft. All authors significantly contributed to the revision of the article and gave final approval for publication.

## Acknowledgements

FM, BR, and JMC gratefully acknowledge the support of the German Centre for Integrative Biodiversity Research (iDiv) Halle-Jena-Leipzig funded by the German Research Foundation (FZT 118).

## Appendix 1: Analytical solution for the persistence probability of aggregated species distributions

In this study, we quantify the abundances and short-term persistence probabilities of species in fragmented landscapes. We use spatial point processes in order to describe species distributions, specifically the Thomas process for aggregated and the Strauss process for regular distributions. In addition, we use fractal maps based on the midpoint displacement algorithm to describe the distribution of habitat in fragmented landscapes. For these fractal maps, we can only assess abundances and persistence probabilities by stochastic simulations. However, for aggregated species distributions in combination with a simplified description of habitat distribution in fragmented landscapes, we derive an analytical solution for the persistence probability as defined in this study. This analytical solution is based on the assumption that the landscape consists of equally sized circular habitat fragments with radius r, which are randomly distributed in the landscape. The number of habitat fragments is called n_F_ and thus the habitat amount equals A_Hab_ = n_F_*π*r^2^ as long as the fragments do not overlap. Accordingly, the proportion of suitable habitat equals A_Hab_/A, while the number of equally sized habitat fragments n_F_ given a fixed habitat amount represents fragmentation *per se*.

As explained in the main text, we assume that all individuals in the habitat fragments survive, while all others outside of the fragments die due to habitat destruction. Accordingly, landscape-level species extinction means that none of the individuals occurs in natural habitat fragments, while landscape-scale persistence means that at least one individual occurs in a habitat fragment (see main text Fig. 1). With these geometric considerations, we can calculate the probabilities of persistence and extinction based on the so-called contact distance distribution

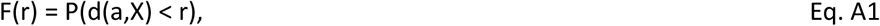

which provides the probability that the distance d between a fragment centred at point a and the nearest individual out of the distribution of individuals (X) prior to landscape change is smaller than the fragment radius r (Fig. S1) (Afshang et al. 2017). The contact distance distribution has been also called empty space function or spherical contact distance distribution (Wiegand and Moloney 2014, Baddeley et al. 2015). Accordingly, F(r) is the probability that at least one individual occurs in the habitat fragment centred at point a, while 1 – F(r) is the probability that there is no individual in this fragment. To quantify the probabilities of extinction or persistence at the landscape scale, we need the probability that there is no individual in any of the n_F_ habitat fragments. Under the assumption that the contact distance distributions F(r) of individual fragments are identical and independent of each other, the probability of extinction is given by:

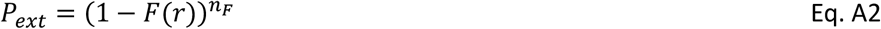

The persistence probability is thus 1 – P_ext_.

In order to link species persistence probabilities to aggregation and abundance, we employ the analytical solution for the contact distance distribution of the Thomas process provided by Afshang et al. (2017):

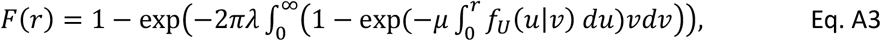

where λ = ρ*μ is the density of the Thomas process and the function f_U_ describes the distribution of individuals’ distances to the fragment centre conditional on a cluster centre’s distance to the fragment centre

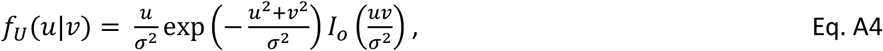

where *I_0_* denotes the modified Bessel function with order zero.

Based on these functions we can calculate the persistence probability of the species as function of its abundance (N = ρ*A*μ), its aggregation (μ and σ), proportion of remaining habitat (n_F_*π*r^2^ /A), and fragmentation *per se* (n_F_).

**Figure A1:**
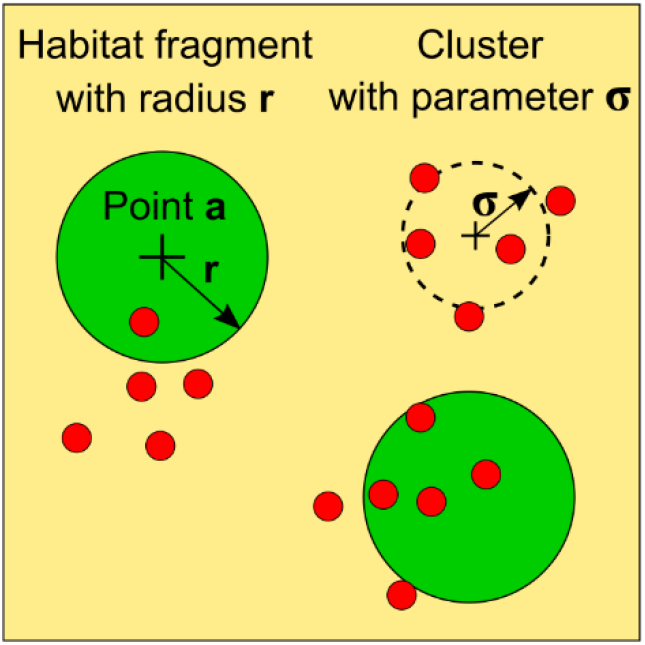
Illustration of the geometric problem to calculate persistence probability. Green area indicates natural habitat and yellow hostile matrix. Each red dot represents an individual of the focal species. Here we show a landscape with two fragments (n_F_ = 2), each with radius r. Accordingly, the total remaining habitat area equals A_Hab_ = n_F_*π*r^2^. The depicted species has an abundance of 15 individuals, with three clusters (n_C_ = 3) and a cluster size parameter of σ. The persistence probability is represented by the probability that at least one individual is located in habitat. We derive an analytical approach to calculate this probability based on the landscape characteristics (number and size of fragments) and species properties (abundance, number and size of clusters).

## Appendix 2: Comparison between the analytical solution and stochastic simulations

The analytical solution described is based on the assertion of identical and independent contact distance distributions for the individual fragments. This assertion implicitly includes several simplifying assumptions. First, we assume that fragments do not overlap, which would introduce non-independence between the F(r) functions of different fragments. Secondly, the analytical solution for the contact distance distribution was derived for infinite areas, while our questions concern finite landscapes, where edge effects might influence the results. Therefore, we checked our analytical results with stochastic spatial simulations in a finite landscape. We simulated species distributions in the same way as described in the main text, but instead of using fractal landscapes, we now used equally sized circular fragments as in the analytical approach. In this case the number of fragments (n_F_) defines fragmentation *per se*, and their total area (n_F_*π*r^2^) habitat amount. We used the so-called hard-core process to simulate the positions of fragment centres. A hard-core process is equal to the Strauss process with inhibition strength parameter γ = 0, which means there is perfect inhibition of points up to inhibition distance R. We set the inhibition distance equal to twice the fragment radius r. In this way, the hard-core process essentially simulates non-overlapping habitat fragments.

We simulated the same full factorial design for species distributions, habitat amount and fragmentation *per se* as described in the main text, but now with landscapes of equally sized circular fragments. The only difference is that fragmentation *per se* is described by varying fragment number from 1 (low fragmentation per se) to 100 (high fragmentation per se) instead of the Hurst factor.

We found close agreement between persistence probabilities from the analytical approach and the stochastic simulations (Figs. A2, A3, A4). However, at higher persistence probabilities the analytical approach sometimes underestimates the simulated values by up to 20%. Specifically the analytical results were biased towards lower persistence probabilities for high habitat amount and a low number of species clusters (Fig. A3, panels in the upper left corner). This bias is caused by the assumption of independence among the contact distributions F(r) of the fragments in the analytical approach (Eq. A2). This assumption is violated in the simulation because of the finite landscape area and the non-overlapping fragments simulated by the hard-core process. However, there was close match between analytical and simulated results for habitat amount below 20% (Fig. A3).

**Figure A2:**
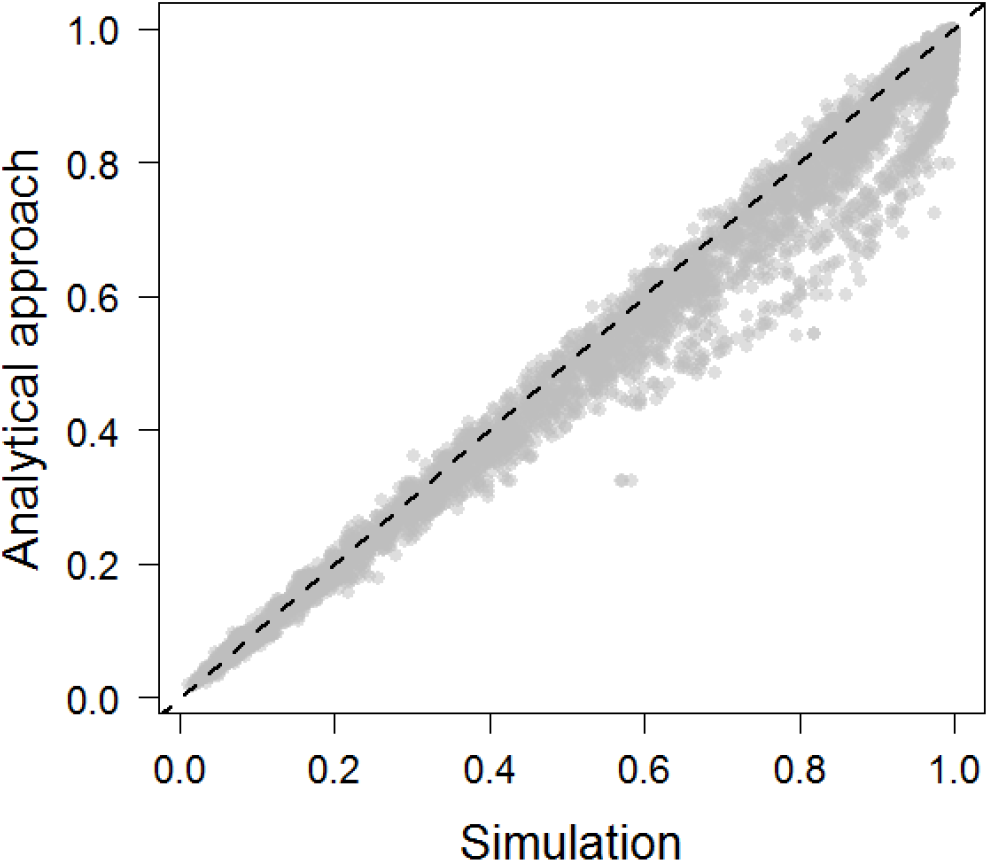
Correlation between persistence probabilities estimated by the stochastic spatial simulation approach and the analytical solution for equally sized circular fragments. This figure shows all used scenarios.

**Figure A3:**
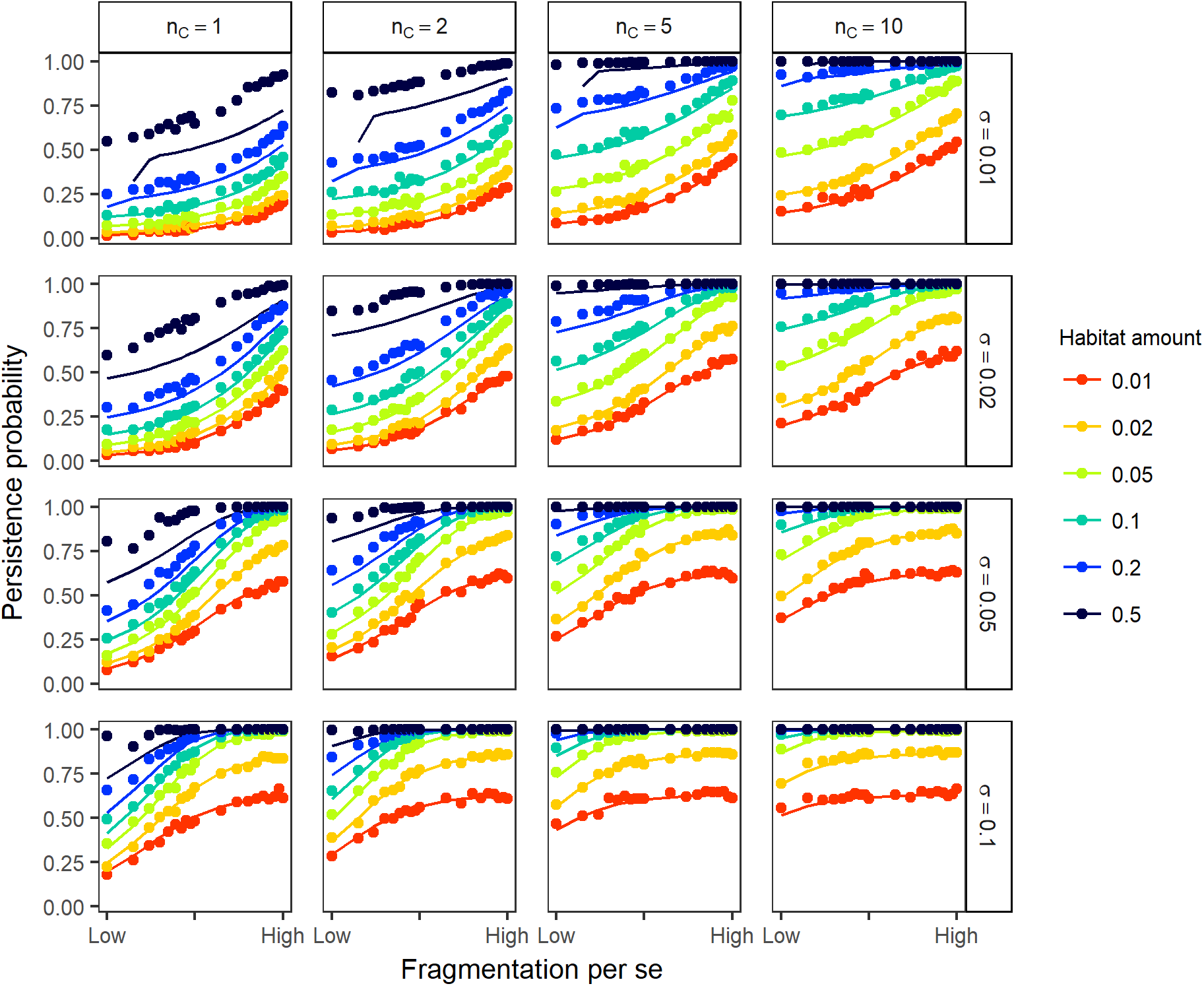
Comparison between persistence probabilities predicted by the stochastic simulation approach and the analytical solution for equally sized circular habitat fragments and aggregated species distributions. Here, the analogous scenarios for species aggregation, habitat amount, and fragmentation *per se* are shown as in Fig. 4 in the main text. The only difference is that fragmentation *per se* is described by the number of fragments (low fragmentation = 1 fragment, high fragmentation = 100 fragments) and not by the Hurst factor. Points show simulation results and lines show results of the analytical approach. The line and point colour indicates the proportion of remaining habitat in the landscape and thus habitat amount. The rows differ in the extent of species clusters as defined by σ, while the columns differ in the number of clusters n_c_. All calculations were done for species with 100 individuals. Simulated persistence probabilities were estimated from 1,000 replicate simulations.

**Figure A4.**
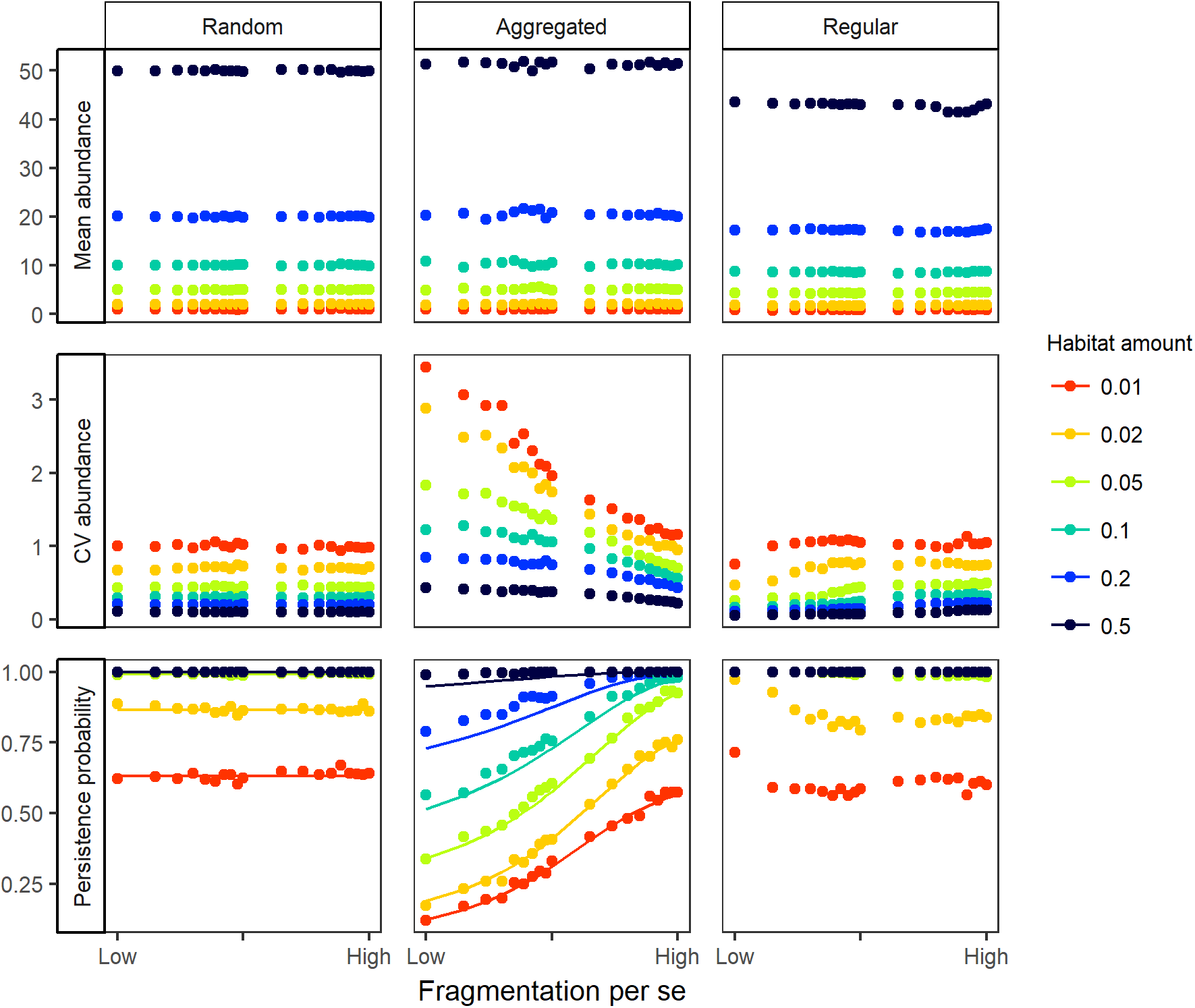
Consequences of habitat amount and fragmentation *per se* for species abundance and persistence with random, aggregated and regular species distributions. In contrast to Fig. 4 in the main text, landscapes with equally sized circular fragments were used here instead of fractal maps. All simulations were conducted with 100 individuals. In the aggregated scenario (middle column) we used a cluster size of σ = 0.02 and a number of clusters n_C_ = 5. In the regular scenario (right column), we used an inhibition strength parameter of γ = 0.01. The rows show the mean abundance (top), coefficient of variation of abundance (= standard deviation/mean) (middle) and the persistence probability (bottom) estimated from 1,000 replicate simulations. Low fragmentation corresponds to 1 fragment and high fragmentation to 100 fragments. The points show simulation results, while the solid lines represent lines from the analytical solution explained in Appendix 1 (note that the analytical approach only provides persistence probabilities and no abundances). The colour of points and lines indicates the habitat amount measured as proportion of habitat in the landscape.

## Appendix 3: Comparison between results for fractal landscapes and landscapes with equally size circular fragments

We also compared results between two different methods for describing the distributions of habitat in fragmented landscapes. We found very close agreement between results for fractal landscapes (Fig. 3 in the main text) and landscapes with equally sized circular fragments (Fig. A4). Only for regular distributions and high habitat amount, we found a lower mean abundance than expected (Fig. A4, upper right panel). This deviation from the theoretical predictions is most likely a result of the highly regular distribution of the circular fragments, which apparently results in an increased probability that individuals are located in the gaps between the equally sized circular fragments compared to the fractal maps.

